# Development of a New Approach Method to Monitor and Modify Caffeine Metabolism Correlated to CYP1A2 Expression

**DOI:** 10.1101/2025.11.27.691042

**Authors:** Xiuyuan Chen, Masaru Miyagi, Gregory P. Tochtrop

**Affiliations:** Department of Chemistry, Case Western Reserve University, Cleveland, Ohio 44106, USA; Department of Pharmacology, Case Western Reserve University, Cleveland, Ohio 44106, USA

**Keywords:** Caffeine, CYP1A2, cell culture, metabolism, LC–MS/MS, Hepatocytes

## Abstract

New approach methodologies (NAMs) that seek to reduce reliance on animal testing require sensitive, mechanism-based assays to accurately predict human-specific metabolic responses. Caffeine, primarily metabolized by cytochrome P450 1A2 (CYP1A2), serves as an ideal probe substrate for evaluating CYP1A2 function. Here, we describe the development of an in vitro platform that combines high-sensitivity triple quadrupole multiple reaction monitoring (MRM) LC-MS analysis of paraxanthine with quantitative reverse-transcription PCR (qPCR) of CYP1A2 expression. Using human hepatocellular carcinoma-derived cell lines (HepG2 and Hep3B), we demonstrate that modulating CYP1A2 with known effectors, sulforaphane (a known CYP1A2 inhibitor), 3-methylcholanthrene (inducer), and galangin (moderate inducer), changes in both paraxanthine accumulation and CYP1A2 mRNA levels can be effectively monitored. The correlations observed between transcriptional responses and metabolic outputs validate paraxanthine as a sensitive readout of CYP1A2 function in these cell lines. Moreover, the assay remains robust across multiple experimental conditions and facilitates insights into enzyme induction or inhibition mechanisms. By providing a straightforward and scalable alternative to animal models, this approach expands the toolbox available for interrogating xenobiotic metabolism, and enzyme regulation. Ultimately, these findings highlight the utility of an integrated cell culture-based system for advancing studies of hepatic enzyme function. This platform enables investigators to readily screen and characterize compounds that influence CYP1A2-mediated metabolism, providing a straightforward, scalable, alternative to animal models.

## Introduction

Efforts to reduce and ultimately eliminate animal testing have gained significant momentum in toxicology, drug discovery, and risk assessment. This movement has led to the emergence of new approach methodologies (NAMs), human-relevant platforms that can supplant traditional in vivo models (1). Within this paradigm, in vitro cell-based assays offer the advantage of potentially better predictive power for human metabolism, while simultaneously adhering to the principles of bioethics and regulatory directives that encourage minimizing animal use. Such approaches capitalize on well-characterized human-derived cell systems, allowing investigators to interrogate key metabolic enzymes without animal experimentation.

Caffeine, a member of the methylxanthine family of natural products (1,3,7-trimethylxanthine), is an ideal substrate for illustrating this type of NAM for the study of hepatic metabolism. It is one of the most widely consumed psychoactive substances globally, inherently contained in or exogenously added to coffee, tea, and various commercial drinks. As such, the metabolism of caffeine has been extensively studied in humans and other species (2). Briefly, caffeine is rapidly absorbed by the gastrointestinal tract and primarily metabolized in the liver by cytochrome P450 1A2 (CYP1A2) (3). Due to active site promiscuity, this enzyme generates three major metabolites with the following approximated relative abundances: 82% paraxanthine (1,7-dimethylxanthine), 11% theobromine (3,7-dimethylxanthine), and 5% theophylline (1,3-dimethylxanthine), see Fig. 1 (4). The *CYP1A2* gene is highly conserved with no common alleles that have been described to impart any important alteration in enzyme activity (5, 6). However, the *CYP1A21F* allele contains a –163C>A mutation in intron 1 that has been shown to influence the inducibility of the gene and display an *in vivo* effect on the magnitude of caffeine metabolism after both smoking (7, 8) and omeprazole treatment (9).

**Figure 1:**
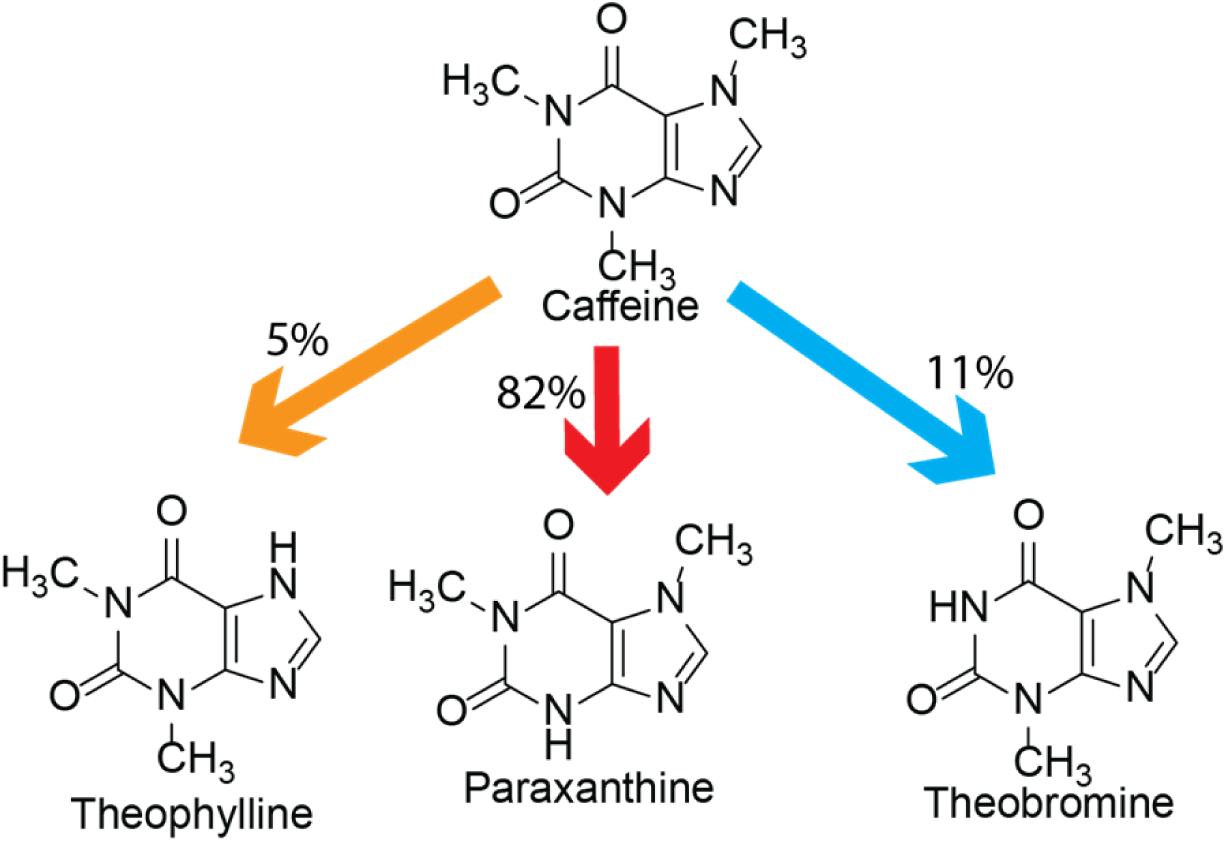
Caffeine is metabolized into three major metabolites by CYP1A2. The relative product distribution of paraxanthine>theobromine>theophylline (82%, 11%, 5%) is due to active site promiscuity in CYP1A2.

We chose cell culture models as they are powerful tools for these types of investigations due to their controlled environment and the ability to manipulate specific variables affecting CYP1A2 expression and activity. In vitro liver models, particularly hepatocellular carcinoma-derived cell lines such as HepG2 and Hep3B, provide valuable platforms for studying hepatic metabolism due to their human origin and stable expression of metabolic enzymes (10). HepG2 cells are derived from a human hepatocellular carcinoma and are frequently used for drug metabolism studies due to their retained liver-specific functions, including the expression of key metabolic enzymes like CYP1A2 (11). Hep3B cells, while also derived from hepatocellular carcinoma, exhibit a distinct metabolic profile that can reveal interindividual variability in drug metabolism (12, 13). The distinct metabolic phenotypes of HepG2 and Hep3B cells can be leveraged to understand interindividual variability and the factors influencing CYP1A2-mediated caffeine metabolism. Further, their relatively stable enzyme expression profiles make them suitable proxies for human liver metabolism, and their use is well-established in vitro studies probing xenobiotic metabolism and drug-drug interactions. Taken together, these features provide a robust experimental framework to investigate not only the basal activity of CYP1A2 but also the extent to which it can be induced or inhibited by exogenous agents.

In this study, we aimed to develop a tractable in vitro method to monitor the inducibility and activity of CYP1A2 in response to bioactive small molecules. Surprisingly, despite the voluminous literature on caffeine metabolism, and the effects of caffeine and its metabolites on various systems, minimal work has been done on caffeine metabolism in culture systems (14). By leveraging the known metabolic pattern of caffeine and its principal metabolites, we first established a high-sensitivity LC separation followed by triple quadrupole mass spectrometric analysis with multiple reaction monitoring (MRM) method for the detection and quantitation of caffeine metabolites, focusing on paraxanthine as a principal readout of CYP1A2 function. We then demonstrated that these metabolites can be reliably measured in HepG2 and Hep3B cultures under various treatment conditions, allowing us to discriminate differences in metabolic rates.

Finally, we evaluated how these metabolic changes correlate with alterations in CYP1A2 mRNA levels, thus linking gene expression to functional activity. This provides a direct correlation between CYP1A2 expression and enzymatic function, offering insight into how various compounds affect caffeine metabolism at both the transcriptional and enzymatic levels. This integrated approach provides a straightforward, in vitro platform for probing CYP1A2 inducibility and serves as a basis for identifying novel modulators of caffeine metabolism or other CYP1A2-mediated processes.

## Results

### Optimization of LC-MS conditions

Numerous analytical methods have been developed and continually refined to facilitate the detection and quantification of caffeine metabolites, including paraxanthine, theobromine, and theophylline, across various biological matrices (15, 16). Early work often relied on gas chromatography-mass spectrometry (GC-MS), leveraging the method’s sensitivity and chromatographic resolution,(15) but their reliance on prior chemical derivatization made sample preparation laborious and added additional steps, which we found to induce additional error and complexity. More recently, studies have increasingly turned to liquid chromatography-mass spectrometry (LC-MS), which offers the key advantage of the ability to forgo derivatization, greatly simplifying workflows. Based on this and influenced by the previous studies by Ma and Mendes (16, 17), we developed a simple, rapid and efficient LC-MS MRM method for simultaneous determination of caffeine and its 2 most abundant metabolites (paraxanthine and theobromine) in cell culture. We chose to forgo analyses of theophylline due to its low relative abundance as a metabolite coupled to the small scale of our experimental cell culture system. To develop the MRM method, we analyzed caffeine, paraxanthine, and theobromine, along with their deuterated internal standards, prepared in water with 0.1% formic acid to a final concentration of 100 μM. Optimal MRM conditions were determined as described in the Figure 2 legend. Based on ion intensity and retention of deuteration patterns, the following precursor/product ion transitions were selected, caffeine 195.1/138.1 and caffeine-d_3_ 198.1/138.0; paraxanthine 180.9/124.0 and paraxanthine-d_6_ 187.1/127.0; theobromine 181.1/138.1 and theobromine-d_6_ 187.1/144.0 (Figure 2A-C). An equimolar mixture of the three analytes was then analyzed by reverse-phase chromatography, and total ion currents (TICs) were monitored to assess chromatographic resolution (Fig. 2D). Caffeine/caffeine-d_3_ eluted at 5.2 min, paraxanthine/paraxanthine-d_6_ eluted at 4.6 min and theobromine-d_6_ eluted at 4.2 min (Fig. 2D). As shown in 2D, theobromine exhibited significantly lower ionization efficiency than paraxanthine or caffeine. Based on this and its approximately eight-fold lower abundance compared to paraxanthine during CYP1A2-mediated demethylation, we chose to forgo theobromine analysis and monitor paraxanthine as a readout of caffeine metabolism.

**Figure 2:**
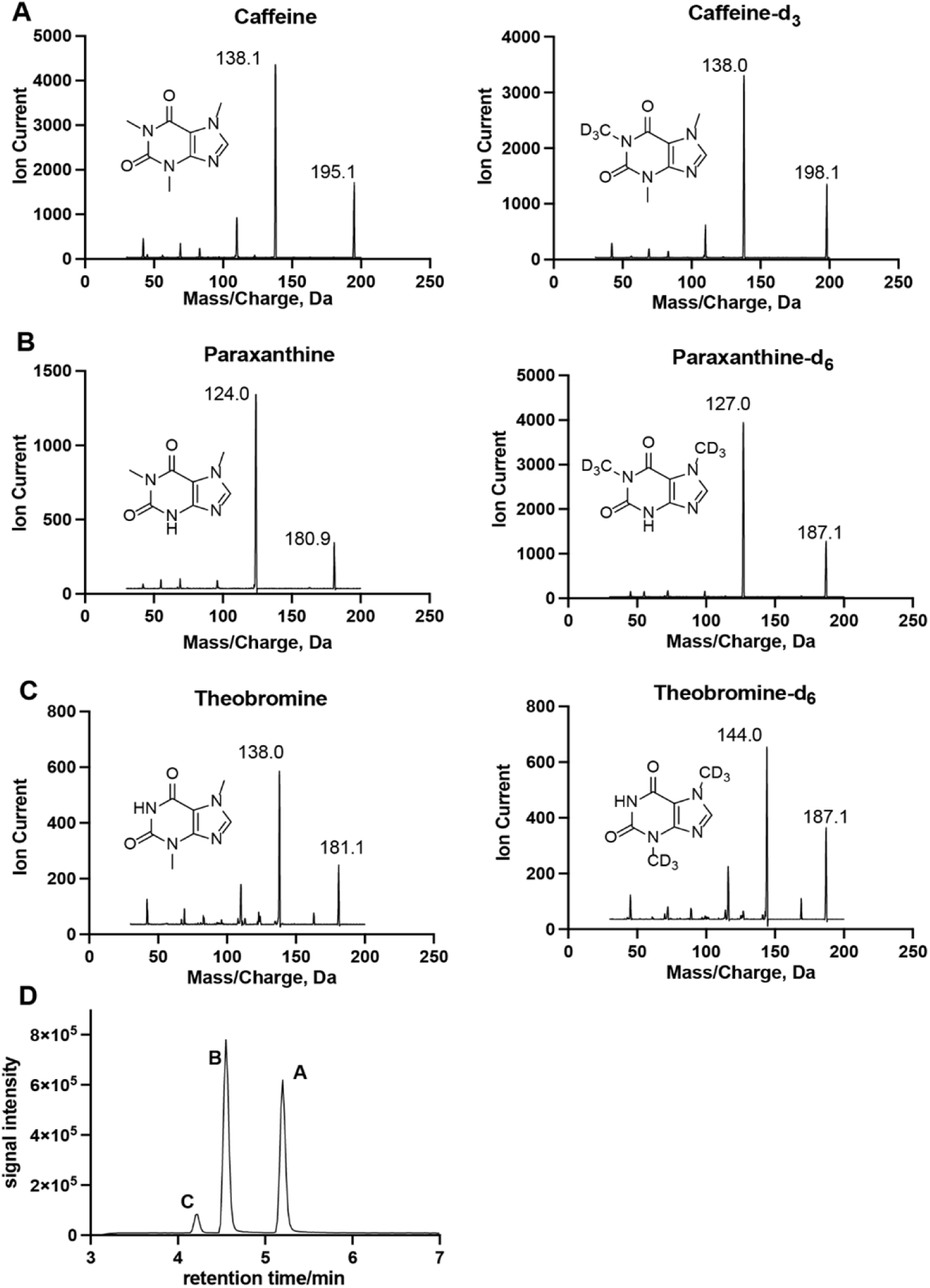
An LC–MS/MS MRM approach to quantifying caffeine metabolites. Quantification of caffeine metabolites is performed through the utilization of deuterated internal standards. **A-C**, show the collision-induced dissociation (CID) product ion spectra of caffeine, paraxanthine, and theobromine, and their internal standards (caffeine-d_3_, paraxanthine-d_6_, and theobromine-d_6_). Spectra were acquired by flow injection analysis (FIA) in positive ion mode. 10 µL of each analyte (10 µM in 0.1% formic acid) was injected into 0.1% formic acid in water-acetonitrile (1:1, v/v) at 200 µL/min, using a fragmentor voltage (FV) of 120 eV and a collision energy (CE) of 20 eV. **D**, MRM total ion chromatogram (TIC) of a 10 µM solution of caffeine (m/z 195.1 → 138.1, FV: 120 eV, CE: 18 eV), paraxanthine (m/z 180.9 → 124.0, FV: 132 eV, CE: 22 eV), and theobromine (m/z 181.1 → 138.1, FV: 120 eV, CE: 22 eV) in 0.1% formic acid in water. Deuterated internal standards were analyzed with the same condition as the standards. MRM conditions, including precursor/product ion selection, fragmentor voltages, and collision energies, were optimized using Agilent MassHunter Optimizer software.

To investigate our ability to extract and quantify paraxanthine in cell culture, we developed the workflow shown in Fig. S7. Linearity and carryover were evaluated by preparing paraxanthine standards prepared in 2 ml of working cell culture media and processing them using the described workflow. Calibration curve for paraxanthine was generated by plotting the peak area ratios of paraxanthine to its deuterated internal standard against the spiked concentrations (0.0125, 0.025, 0.05, 0.1, 0.2 and 0.4 µM). This calibration curve covers the whole experimental working range with acceptable linearity (R^2^ >0.99, Fig. S1). Carryover was assessed by analyzing 5 blank samples after each injection of standard samples, all of which showed < 0.1% residual paraxanthine.

To establish a robust in vitro platform for investigating caffeine metabolism, we began by assessing the tolerance of HepG2 and Hep3B cells to caffeine using an MTT viability assay, which revealed that concentrations up to 1 mM had no significant effect on viability (Fig. S2). Our initial challenge was to determine whether changes in paraxanthine concentrations could be determined over time in culture. To our surprise, we discovered that all tested commercial sources of caffeine contained approximately 0.1% paraxanthine, generating an issue that based on variability in paraxanthine impurity concentration in different commercial sources of caffeine, the overall production of paraxanthine could not be precisely detected, only changes from the initial concentration. This necessitated the purification of caffeine by preparative HPLC as described in materials and methods. After this initial hurdle, we were able to detect accumulation of paraxanthine as a function of time (Fig. S6) with samples collected at 0, 2, 4, 6, and 24 hours. The resulting concentration profiles were used to calculate accumulation rates via linear trendlines (Table 1, Fig. S6). A blank group, containing 1 mM caffeine but no cells, was included to account for trace paraxanthine in the purified caffeine as well as any concentration changes due to media evaporation, thus providing an accurate baseline for subsequent comparisons.

**Table 1.** Paraxanthine accumulation rates in HepG2 and Hep3B cells under different treatments. The accumulation rate (nM/h) was determined from the difference in paraxanthine concentration at 0 and 24 hours. “Blank” groups received 1 mM caffeine and DMSO (vehicle), whereas “Treat” groups were exposed to the same caffeine concentration plus 10 µM D,L-sulforaphane, 0.5 µM 3-methylcholanthrene, or 10 µM galangin. Data are presented as mean ± SD (n = 3).

| Treatment | Cell Type | Group | Paraxanthine<br>accumulation in 24h<br>(nM/h) |
| --- | --- | --- | --- |
| D, L-Sulforaphane | HepG2 | Blank | 3.0 ± 0.5 |
|  |  | Treat | 1.4 ± 0.4 |
|  | Hep3B | Blank | 0.5 ± 0.2 |
|  |  | Treat | 0.4 ± 0.1 |
| 3-Methylcholanthrene | HepG2 | Blank | 1.5 ± 0.9 |
|  |  | Treat | 49.7 ± 4.4 |
|  | Hep3B | Blank | 0.5 ± 0.2 |
|  |  | Treat | 5.2 ± 1.6 |
| Galangin | HepG2 | Blank | 0.7 ± 0.1 |
|  |  | Treat | 2.8 ± 0.3 |
|  | Hep3B | Blank | 0.5 ± 0.2 |
|  |  | Treat | 1.6 ± 0.2 |

### Investigations of bioactive small molecule treatments on HepG2/Hep3B caffeine metabolic rate by monitoring paraxanthine concentration changes in cell culture

We selected sulforaphane, 3-methylcholanthrene (3-MC), and galangin because each compound modulates CYP1A2 via distinct mechanisms, thereby providing a broad test of our caffeine-metabolism assay. Sulforaphane is known to inhibit CYP1A2 activity, making it a representative suppressor of metabolism (18). In contrast, 3-MC, a classical aryl hydrocarbon receptor (AHR) agonist, robustly induces CYP1A2 expression (19). Galangin, another AHR activator, offers a more moderate but still measurable inductive effect (17, 19). Together, these three diverse modulators allowed us to explore both the inhibitory and inducible ranges of CYP1A2 function within our cell-culture model. Next, we verified the compatibility (toxicity) of these compounds in our cell culture system (Fig. S3–S5). Specifically, control cells were incubated with 1 mM caffeine alone, whereas treatment groups received 1 mM caffeine plus sulforaphane, galangin, or 3-MC at the indicated concentrations based on the MTT toxicity data.

We then examined the ability of these compounds to alter caffeine metabolism in our cell culture model. Compared to the control group, sulforaphane decreased paraxanthine accumulation to 63% of the control (p<0.05) in HepG2 cells after 24 h incubation (Fig. 3A) and decreased paraxanthine accumulation to 87% of control (p>0.05) on Hep3B cells after 24 h incubation (Fig. 3B). 3-MC significantly increased paraxanthine accumulation by 16.7-fold (p<0.0001) on HepG2 cells (Fig. 3E) and by 4.43-fold (p<0.05) on Hep3B after 24 h incubation (Fig. 3F). Galangin significantly increased paraxanthine accumulation by 2.44-fold (p<0.001) on HepG2 cells (Figure 3C) and by 2.04-fold (p<0.01) on Hep3B (Figure 3D) after 24 h incubation.

**Figure 3.**
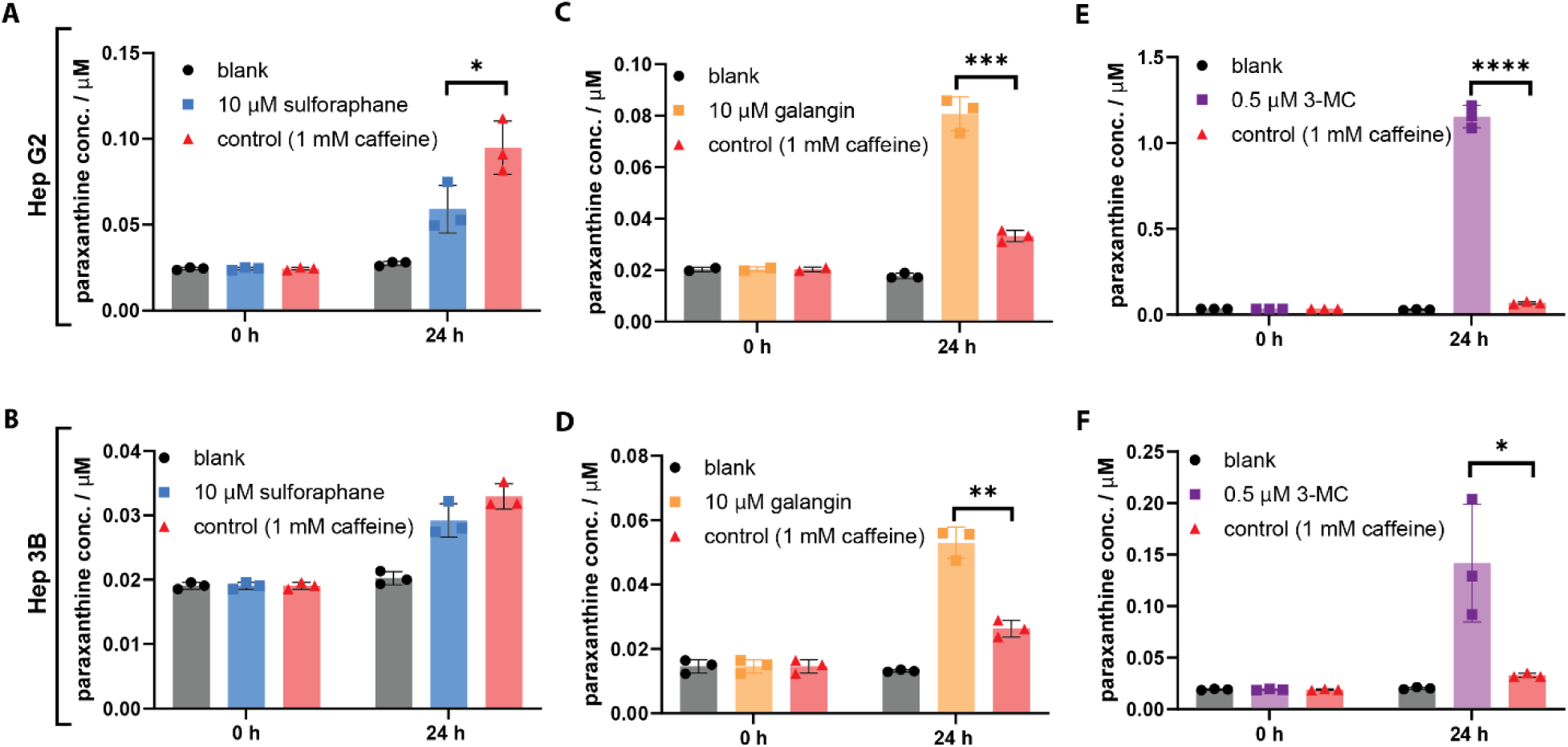
Modulation of paraxanthine accumulation by known CYP1A2 effectors in HepG2 and Hep3B cells. Paraxanthine concentration was quantified by LC–MRM. Data are shown as mean ± SD from three independent experiments. **A-B,** Sulforaphane reduced paraxanthine accumulation in HepG2 and Hep3B cells. **C-D**, Galangin increased paraxanthine accumulation in both cell lines. **E-F**, 3-Methylcholanthrene (3-MC) also increased paraxanthine accumulation in both cell lines. The results of t-test are reported as *p < 0.05, **p < 0.01, ***p < 0.001, ****p < 0.0001.

### Effects of different treatments on HepG2/Hep3B CYP1A2 mRNA-Expression levels

Given that caffeine’s primary metabolism is carried out by CYP1A2, we wanted to determine if the observed perturbations in paraxanthine accumulation were correlated to CYP1A2 expression. Accordingly, we used quantitative reverse-transcription PCR (qPCR) to measure CYP1A2 mRNA levels in HepG2 and Hep3B cells. Given that caffeine itself has been reported to induce CYP1A2 expression in rat hepatocytes(20), the first set of experiments was focused on understanding the effect of 1mM caffeine on baseline CYP1A2 expression. As shown in Fig. 4A and 4B, qPCR revealed that 1 mM caffeine significantly upregulated CYP1A2 mRNA levels in both cell lines, increasing expression by 2.05-fold in HepG2 (p<0.0001) and by 1.85-fold in Hep3B (p<0.05), relative to vehicle-only controls (0.1% DMSO with no caffeine). Based on this observation, cells treated with 1 mM caffeine alone were used as the “control” in subsequent experiments to compare the additional effects of each test compound. We next assessed the impact of sulforaphane on CYP1A2 expression. In HepG2 cells, sulforaphane significantly reduced CYP1A2 mRNA levels to 0.54-fold (p<0.0001) compared to the caffeine-only control (Fig. 4C). In Hep3B cells, sulforaphane lowered CYP1A2 mRNA by 0.72-fold, although this reduction was not statistically significant (p>0.05) (Figure 4D). 3-MC produced a robust induction of CYP1A2 in both cell lines. Relative to caffeine-only controls, 3-MC increased CYP1A2 expression by 2.92-fold (p<0.01) in HepG2 (Fig. 4E) and by 3.81-fold (p<0.05) in Hep3B (Fig. 4F). These data confirm the potent ability of 3-MC to stimulate CYP1A2 transcription. Finally, galangin was evaluated for its effect on CYP1A2 expression. In HepG2 cells, CYP1A2 mRNA increased by 4.77-fold (p>0.05, ns) compared to the caffeine-only control (Fig. 4G). In Hep3B cells, galangin triggered a 1.84-fold rise (p<0.01), indicative of a moderate but statistically significant increase in CYP1A2 expression (Fig. 4H). Overall, these data establish that each of the tested modulators, sulforaphane, 3-MC, and galangin, can differentially affect CYP1A2 expression in human liver cell lines. Moreover, the observed changes in mRNA levels are consistent with the trends in paraxanthine accumulation (Fig. 3), suggesting that transcriptional regulation by these compounds translates into functional CYP1A2 activity.

**Figure 4.**
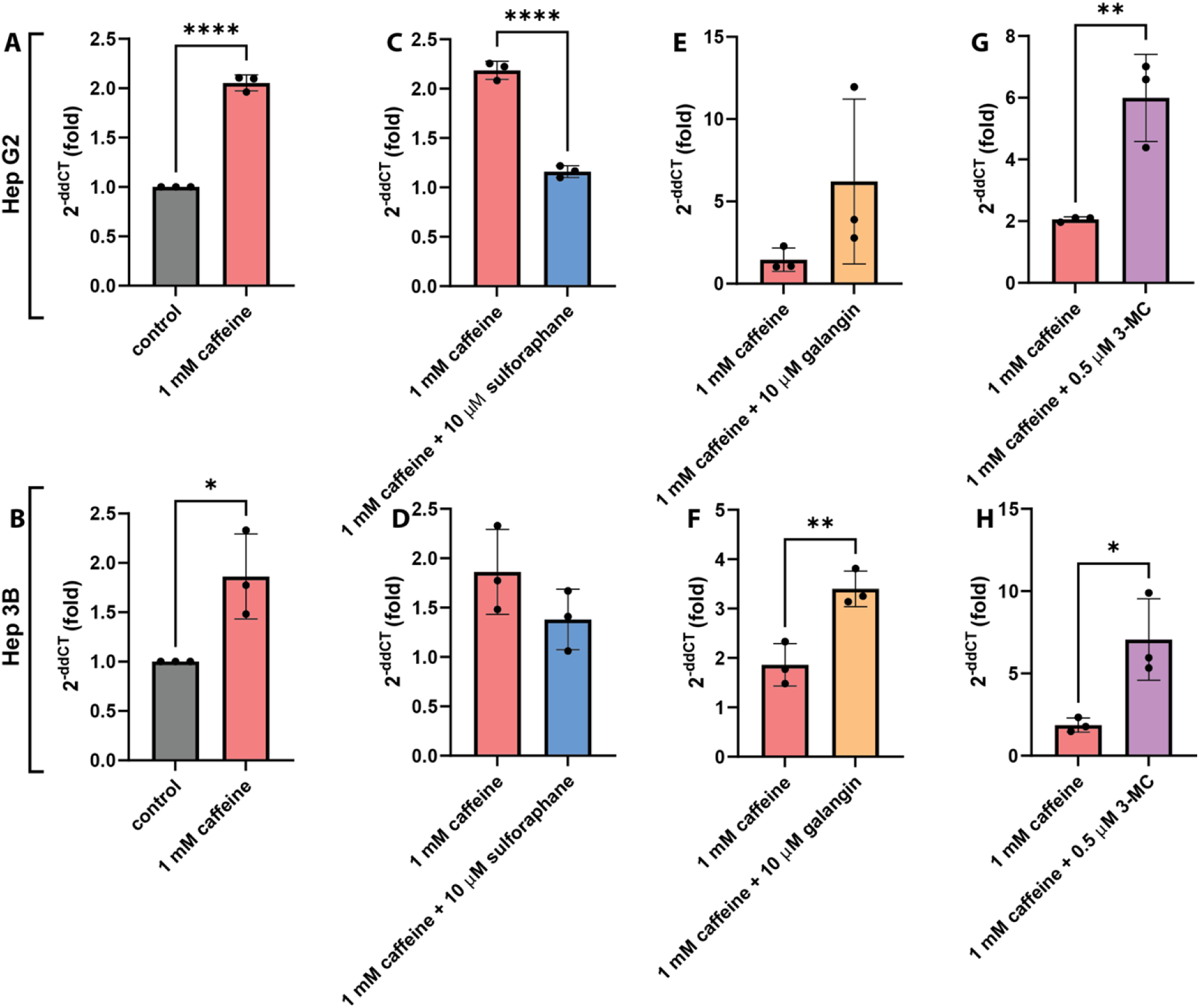
qPCR measurement of CYP1A2 mRNA levels in HepG2 (top panels) and Hep3B (bottom panels) cells after 24-hour treatments with **A-B**, caffeine and caffeine along with **C-D,** D,L-sulforaphane, **E-F**, 3-methylcholanthrene, or **G-H**, galangin at the indicated concentrations. Control cells received the vehicle only. Data are normalized to a housekeeping gene and expressed as the mean ± SD (n = 3) from three independent experiments. The results of t-test are reported as *p < 0.05, **p < 0.01, ***p < 0.001, ****p < 0.0001.

## Discussion

In this study, we developed and validated an in vitro method to monitor caffeine metabolism in hepatocellular carcinoma-derived cell lines (HepG2 and Hep3B) by specifically measuring paraxanthine accumulation as a sensitive readout of CYP1A2 activity. As a NAM, this integrated platform capitalizes on human-relevant cellular models to investigate xenobiotic metabolism without the ethical and logistical complexities associated with animal testing. By combining a straightforward LC–MRM analysis with qPCR, we provide a robust and reproducible alternative for interrogating CYP1A2 modulation, thereby aligning with current regulatory and scientific priorities to reduce, refine, and replace in vivo models. Within the results comprising this report, we highlight three key advances.

### Mechanistic concordance between transcription and catalysis

Across two genetically distinct liver-derived lines, sulforaphane, galangin and 3-MC produced parallel, direction-consistent changes in *CYP1A2* mRNA and paraxanthine output. This tight correlation confirms that measuring transcript levels could be used as a quick screening tool to predict functional activity, which is especially useful when the mass spectrometry assay is more labor-intensive. In our study, we indeed see that a significant induction or inhibition at the mRNA level translated into a proportional change in metabolite output.

### Assay robustness and analytical sensitivity

The LC–MRM workflow achieved low-nanomolar detection in culture medium with minimal carry-over, enabling rate measurements from a single 24 h time-course. Importantly, preparative HPLC removed trace paraxanthine contamination present in commercial caffeine lots, an underappreciated confounder that would otherwise obscure subtle metabolic shifts. The resulting dynamic range captured a 17-fold induction (3-MC in HepG2) and a 40% inhibition (sulforaphane), demonstrating suitability for both strong and modest modulators.

### Alignment with regulatory drivers toward animal-free testing

Moreover, as the field of metabolism and toxicology research continues to move away from reliance on animal models and towards NAMs, the development of robust, high-throughput in vitro systems becomes paramount. his combined LC–MS/MS plus qPCR platform offers an attractive in vitro alternative to in vivo studies for CYP1A2 modulation, helping reduce reliance on animal experiments. The consistent performance of this method in both HepG2 and Hep3B cells further underscores its potential utility as a screening platform for CYP1A2 modulators, drug–drug interactions, nutraceutical impacts, or environmental toxicants. Looking forward, these findings highlight the promise of cell-based assays for advancing personalized medicine and refining our understanding of how complex metabolic enzymes like CYP1A2 respond to a variety of chemical stressors.

While caffeine clearance has long been exploited in vivo to phenotype CYP1A2, in vitro methods have lagged. Previous culture studies focused on primary hepatocytes, which are scarce, variable and rapidly lose enzyme expression (21). By contrast, HepG2 and Hep3B cells maintain stable growth and permit genetic or pharmacological manipulation; Hepatoma lines like HepG2/Hep3B generally have lower absolute CYP enzyme expression than primary hepatocytes, which is why a highly sensitive method (like LC–MS/MS) was essential. Our data now show they can also reveal CYP1A2 modulation when paired with a sufficiently sensitive analytical readout.

In conclusion, our findings illustrate how a cell-based assay centered on caffeine metabolism and paraxanthine production can serve as a practical NAM. By providing a clear link between transcriptional regulation and functional enzymatic output, this integrated system offers a reliable means to screen for novel CYP1A2 modulators. It deepens our understanding of xenobiotic metabolism and helps advance the broader shift toward non-animal alternatives in toxicology and pharmacology.

## Materials and methods

### Reagents

Solvents were purchased from Fisher Scientific. Caffeine, caffeine-d_3_, paraxanthine, paraxanthine-d_6_, theobromine and theobromine-d_6_ were analytical standards grade (purity >99%) and purchased from Cayman Chemical. D, L-Sulforaphane, 3-Methylcholanthrene (3-MC), Galangin and DMSO were purchased from Sigma-Aldrich. Caffeine utilized for cell culture experiments was further purified by preparative reverse phase HPLC on a Shimadzu LC-20AP using an Agilent Prep-C18 column (PN-410910-102 21.2 × 250 mm). Human HCC cell lines Hep3B (HB-8064) and HepG2 (HB-8065) were purchased from ATCC (Manassas, VA, USA). CyQUANT MTT Cell Viability Assay and High-Capacity cDNA Reverse Transcription Kit were purchased from ThermoFisher Scientific. RNeasy Mini Kit was purchased from QIAGEN. qRT-PCR was performed using QuantStudio 3 Real-Time PCR System from Applied Biosystems and TaqMan® Gene expression assays (Applied Biosystems). The following TaqMan probes (Applied Biosystems) were used: human CYP1A2 (Hs00167927_m1), human GAPDH (Hs02786624_g1).

### Cell culture experimental design

HepG2 and Hep3B cells were seeded in 6 well plates at a density of 1 × 10^6^ cells/well. Minimum Essential Media (MEM, Gibco, USA) which contains 10% Fetal Bovine Serum (FBS, gibco, USA) and 1% penicillin-streptomycin (Cytiva, USA) was used to maintain the cell growth. All cells were grown in the medium overnight to settle down before treatment.

### HPLC method for caffeine and its major metabolites separation

Analytes were separated by reverse-phase chromatography (Kinetex 2.6 µm Polar C18 100 Å, LC Column 100 × 2.1 mm) on an Agilent 1290 HPLC system. Mobile phase A and B were 0.1% formic acid in water and 0.1% formic acid in acetonitrile. LC program for separation: 0–50 % of B (0–8 min), 50-80 % of B (8-10 min), 80-0% of B (10-15 min) and flow rate was 0.2 ml/min.

### Monitoring CYP1A2 expression in HepG2/Hep3B cells with different treatments

Control cells were incubated with the same growth media plus 1% of DMSO. For experimental cells, 2 ml of 1 mM caffeine growth media was used in each well, and 1 mM caffeine growth media was made by dissolving solid caffeine powder into growth media. 20 µl of either DMSO, 50 µM 3-MC in DMSO, 1 mM sulforaphane in DMSO or 1 mM galangin in DMSO was added into different experimental cell groups. Experimental groups received 1 mM caffeine + modulator (final concentrations: 10 µM sulforaphane, 0.5 µM 3-MC, or 10 µM galangin, with DMSO vehicle at 1% v/v). Concentration of treatments was determined by MTT cell toxicity assay in Figure S2-5. After 24 h incubation, cells were washed with PBS and then treated with trypsin to collect the cell pellets. Total RNA was extracted from cell pellets with RNeasy Mini Kit (QIAGEN, USA). RNA quality was determined by the 260/280 nm absorbance ratio and concentration was obtained by Nanodrop Microvolume Spectrophotometers (Thermo Scientific, USA). 5 µg of RNA from each sample was reverse transcribed to cDNA by using High-Capacity cDNA Reverse Transcription Kit (ThermoFisher Scientific, USA). qRT-PCR was performed by QuantStudio 3 Real-Time PCR System using ΔΔCT experiment type and fast running mode.

GAPDH (Thermo Fisher assay ID Hs02786624_g1) served as the internal reference gene and CYP1A2 (Thermo Fisher assay ID Hs00167927_m1) was the target gene. Thermal cycling was performed as follows: 50 °C for 2 min (reverse transcription), 95 °C for 2 min (initial denaturation), then 40 cycles of 95 °C for 1 s and 60 °C for 20 s. The relative mRNA-expression was calculated by 2^-ΔΔCT^ method.

### Time course experiments of monitoring paraxanthine concentration changes in cell culture

Cells were pre-treated for 24 h with the respective modulators prior to the time-course experiment. Control cells received 1% DMSO (vehicle) during this pre-treatment, whereas experimental groups received either 0.5 µM 3-MC or 10 µM sulforaphane (with DMSO as vehicle at 1%). After 24 h pre-treatment, the media were replaced with fresh growth media containing 1 mM caffeine (purity >99%, further purified via preparative HPLC to remove trace paraxanthine). The same concentrations of 3-MC or sulforaphane (or DMSO for controls) were present during the time-course as well. After 24 h of incubation with treatments, the old media were removed and replaced with fresh 2 ml growth media which contains 1 mM caffeine. Same concentrations of treatments were also added during the time course incubations. Samples were collected at the time of 0, 2, 4, 6, 24 h after adding growth media with caffeine and treatments.

### Cell metabolites extraction and LC-MS sample preparation

3 mL of methanol containing 0.13 µM paraxanthine-d6 internal standard was added to each well. Cells were scraped to detach them, and the entire well contents were transferred to a 5 mL tube. Samples were then centrifuged at 14,000×g for 10 min to precipitate debris. 250 µl of supernatant was collected into a new microcentrifuge tube and then dried with a speedvac vacuum concentrator. Samples could then be stored at – 80 °C for up to a week. For LC-MS sample preparation, 100 µl water which contains 0.1% formic acid was added into the dry cell extraction sample. After vortexing to ensure the extract was fully dissolved, 10 µL of the solution was injected into the LC–MS

## Acknowledgements

The authors gratefully acknowledge the financial support provided by the J.M. Smucker Corporation, which made this research possible. The content and conclusions presented herein are those of the authors and do not necessarily represent the views or positions of the J.M. Smucker Corporation.

